# Integrated phospho-proteogenomic and single-cell transcriptomic analysis of meningiomas establishes robust subtyping and reveals subtype-specific immune invasion

**DOI:** 10.1101/2021.05.11.443369

**Authors:** Christina Blume, Helin Dogan, Lisa Schweizer, Matthieu Peyre, Sophia Doll, Daniel Picard, Roman Sankowski, Volker Hovestadt, Konstantin Okonechnikov, Philipp Sievers, Areeba Patel, David Reuss, Mirco Friedrich, Damian Stichel, Daniel Schrimpf, Katja Beck, Hans-Georg Wirsching, Gerhard Jungwirth, C Oliver Hanemann, Katrin Lamszus, Manfred Westphal, Nima Etminan, Andreas Unterberg, Christian Mawrin, Marc Remke, Olivier Ayrault, Peter Lichter, Stefan M Pfister, Guido Reifenberger, Michael Platten, Till Milde, David TW Jones, Rachel Grossmann, Zvi Ram, Miriam Ratliff, Christel Herold-Mende, Jan-Philipp Mallm, Marian C Neidert, Wolfgang Wick, Marco Prinz, Michael Weller, Matthias Mann, Michel Kalamarides, Andreas von Deimling, Matthias Schlesner, Felix Sahm

**Affiliations:** Dept. of Neuropathology, University Hospital Heidelberg and CCU Neuropathology, German Cancer Research Center; German Consortium for Translational Cancer Research, Heidelberg, Germany; Department of Proteomics and Signal Transduction, Max Planck Institute of Biochemistry, Martinsried, Germany; NNF Center for Protein Research, Faculty of Health Sciences, University of Copenhagen, Copenhagen, Denmark; OmicEra Diagnostics GmbH, Planegg, Germany; Sorbonne Université and Department of Neurosurgery, Pitié Salpêtrière Hospital, Paris, France; Dept. of Pediatric Neuro-Oncogenomics, German Cancer Consortium and German Cancer Research Center, Düsseldorf, Germany; Department of Neuropathology, Medical Faculty, University Hospital Düsseldorf, Düsseldorf, Germany; Institute of Neuropathology, Faculty of Medicine, University of Freiburg, Freiburg, Germany; Center for Basics in NeuroModulation (NeuroModulBasics), Faculty of Medicine, University of Freiburg, Freiburg, Germany, Signalling Research Centres BIOSS and CIBSS, University of Freiburg, Freiburg, Germany, Pediatric Oncology, Dana Faber Cander Institute, Boston, USA; German Cancer Consortium, Clinical Cooperation Unit (CCU) Neuroimmunology and Brain Tumor Immunology, German Cancer Research Center, Heidelberg, Germany; Department of Neurology, University Hospital and Medical Faculty Mannheim, Mannheim, Germany; Dept. of Translational Medical Oncology, National Center for TumorDiseases (NCT), Heidelberg, Germany; Division of Molecular Genetics, German Cancer Research Center, Heidelberg, Germany; Department of Neurology and Brain Tumour Centre, Cancer Centre Zürich, University Hospital and University of Zürich, Zürich, Switzerland; Dept. of Neurosurgery, University Hospital Heidelberg, Heidelberg, Germany; Faculty of Health, Peninsula Medical School, University of Plymouth, Plymouth, UK; Dept. of Neurosurgery, University Hospital Hamburg; Dept. of Neurosurgery, University Hospital Mannheim, Mannheim, Germany; Dept. of Neuropathology, University Hospital Magdeburg, Institut Curie, PSL Research University, CNRS UMR, INSERM, Orsay, France; Université Paris Sud, Université Paris-Saclay, CNRS UMR 3347, INSERM U1021, Orsay, France; Hopp Children’s Cancer Center, Heidelberg, Germany; Division of Pediatric Neurooncology, German Cancer Consortium and German Cancer Research Center, Heidelberg, Germany; Department of Pediatric Hematology and Oncology, Heidelberg University Hospital, Heidelberg, Germany; Dept. of Neuropathology, University Hospital Düsseldorf, Düsseldorf, Germany; Pediatric Glioma Research Group, German Cancer Research Center, Heidelberg, Germany; Department of Neurosurgery, Tel Aviv Medical Center, Tel Aviv, Israel, and Sackler School of Medicine, Tel Aviv University, Tel Aviv, Israel; Division of Chromatin Networks, German Cancer Research Center and Bioquant, Heidelberg, Germany; Department of Neurosurgery and Clinical Neuroscience Center, University Hospital and University of Zurich, Zurich, Switzerland; Department of Neurosurgery, Kantonsspital St. Gallen and Medical School St. Gallen, St. Gallen, Switzerland; Clinical Cooperation Unit Neurooncology, German Consortium for Translational Cancer Research, German Cancer Research Center, Heidelberg, Germany; Department of Neurology and Neurooncology Program, National Center for Tumor Diseases, Heidelberg University Hospital, Heidelberg, Germany; Biomedical Informatics, Data Mining and Data Analytics, Faculty of Applied Computer Science and Medical Faculty, University of Augsburg, Augsburg, Germany

## Abstract

Meningiomas are the most frequent primary intracranial tumors. They can follow a wide clinical spectrum from benign to highly aggressive clinical course. No specific therapy exists for refractory cases or cases not amenable to resection and radiotherapy. Identification of risk of recurrence and malignant transformation for the individual patients is challenging. However, promising molecular markers and prognostic subgrouping by DNA methylation are emerging. Still, the biological underpinnings of these diagnostic subgroups are elusive, and, consequently, no novel therapeutic options arise thereof. Here we establish robust subgroups across the full landscape of meningiomas, consistent through DNA methylation, mutations, the transcriptomic, proteomic and phospho-proteomic level. Pronounced proliferative stress and DNA damage repair signals in malignant cells and in clusters exclusive to recurrent tumors are in line with their higher mitotic activity, but also provide an explanation for the accumulation of genomic instability in anaplastic meningiomas. Although homozygous deletion of *CDKN2A/B* is a diagnostic marker of high-grade meningioma, the expression of its gene product increased from low to non-deleted high-grade cases. Differences between subgroups in lymphocyte and myeloid cell infiltration, representing a majority of tumor mass in low-grade NF2 tumors, could be assigned to cluster-specific interaction with tumor cells. Activation to a more proinflammatory phenotype and decreased infiltration of myeloid cells in high-grade cases correlated with lower expression of *CSF1*, located on chromosome arm 1p, whose deletion is known as prognostic marker, with no proposed mechanism before. Our results demonstrate a robust molecular subclassification of a tumor type across multiple layers, provide insight into heterogeneous growth dynamics despite shared pathognomonic mutations, and highlight immune infiltration modulation as a novel target for meningioma therapy.

## INTRODUCTION

Meningiomas are the most frequent primary Central Nervous System (CNS) tumors^1^. They arise from the arachnoidal layer of the meninges and can follow highly divergent clinical courses. While some meningiomas are incidentally detected at imaging or autopsy, most cause symptoms that can only be relieved by resection. Some of these recur despite multiple surgeries and radiation therapy, and patients succumb to the disease^2^. The challenges to translational meningioma research thus comprise both the reliable identification of prognostic factors for the individual patient, and the development of novel treatment concepts for tumors not controlled by surgery and radiation alone.

Besides traditional histopathological evaluation, molecular parameters have been proposed as risk predictors. Several studies have elucidated a dichotomy in the molecular meningioma landscape, with *NF2* altered cases accounting for a about two third on the one, and cases with a variety of other mutations, mostly *AKT1, KLF4, TRAF7* and *SMO*, on the other hand^3–7^. Strikingly, *NF2* mutations can occur across the entire spectrum of clinical manifestations, from benign to highly aggressive, whereas the others are restricted to low-grade cases^7,8^. Thus, patients with meningiomas harboring a targetable alteration such as in *AKT1* or *SMO* are typically not in need for any adjuvant treatment thanks to the benign nature of the tumor^7,9^. In turn, presence of an *NF2* alteration neither provides an established target, nor reliably informs about the risk of reccurrence - since *NF2* mutations are found across the entire spectrum of malignancy.

In order to identify prognostic subgroups of meningioma that overcome the limitations of stratification by mutation or the necessarily subjective and sampling-dependent morphological assessment, we devised an epigenetic classification of meningiomas based on DNA methylation-derived subgroups^8^. Stratification for these subgroups has higher predictive power than the grading criteria of the WHO classification of brain tumors. Six methylation classes (MC) were delineated, three of which display a benign outcome (MC ben-1, 2, 3), two with intermediate course (MC int-A, B), and one with highly aggressive growth (MC mal). These six MCs correlate not only with outcome, but also with other molecular characteristics: While ben-2 encompasses cases with *AKT1, SMO, KLF4, TRAF7* mutations, the *NF2* mutant cases are found in the other five MCs. In line with the data on copy-number variations (CNVs), there is a benign MC with only 22q deletion, the arm on which *NF2* is located, and the number of CNVs increases in the more aggressive MCs int-A, B and mal^8,10^.

Although this epigenetic classification is highly valuable for risk prediction, it still leaves the underlying biological mechanisms unexplained that transform a low-grade to a high-grade meningioma, and why only the *NF2*-mutant cases seem to be susceptible to this transformation.

Here, we set out to identify the decisive steps of meningioma progression on RNA level in bulk tissue samples, proceed with single-cell resolution to identify common and distinct subpopulations in meningiomas across the stages of malignancy, and correlate these findings with their translational effects on the proteomic level.

## RESULTS

### Subgroups are consistently recapitulated on multiple levels

To characterize meningioma subtypes based on the transcriptional as well as on the proteome level, RNA sequencing (RNA-Seq) and proteomics data was generated for a cohort comprising 44 meningioma samples, with both data types being available for 40 of them. The sample set comprised tumors of WHO grades 1, 2, and 3, and of MCs ben-1, ben-2, and mal (Fig. 1A), representing *NF2* low grade (ben-1), *NF2* high grade (mal) and non- *NF2* cases (ben-2). As the MC classification is based purely on methylation data, we sought to investigate whether the separation observed there can be recapitulated on mRNA and protein expression levels. To this end, similarity matrices were generated, in which each pair of samples was allocated a score based on similarity in their expression profiles for RNA-Seq and proteomics data individually. In a next step, a similarity network fusion (SNF) analysis was conducted combining both data modalities. In the SNF, all three MCs clearly separated (Fig. 1B). Thus, the individual MCs can be recapitulated by the combined information of mRNA and protein expression profiles.

**Figure 1.**
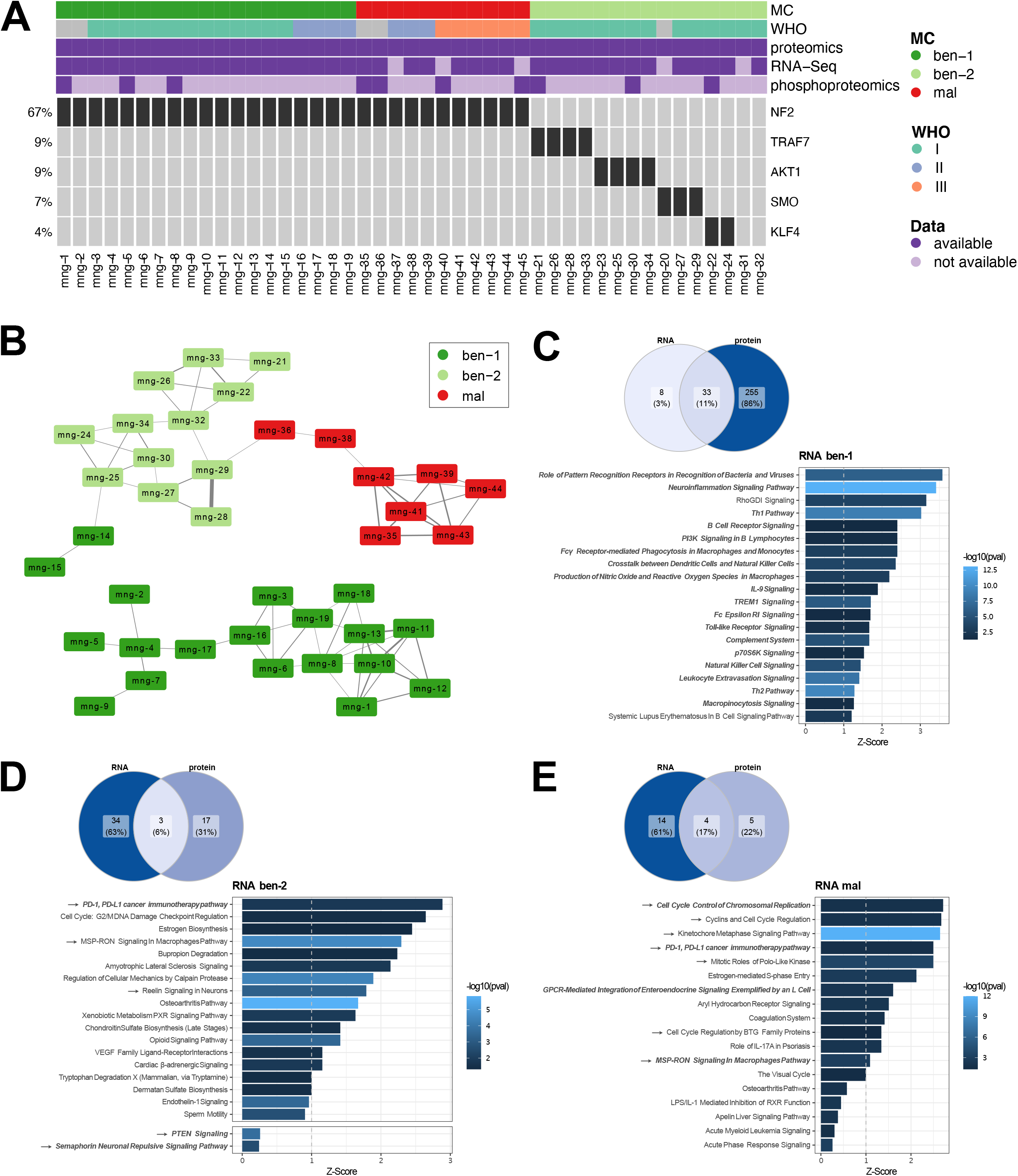
**A** Meningioma dataset used for bulk RNA-Seq, proteomics, and phosphoproteomics analyses with annotation of WHO grade, MC and mutation status for each tumor. **B** SNF graph deducted from bulk RNA-Seq and proteomics data. Each node represents a sample with colors indicating the respective MC. Edges between nodes indicate similarity between the sample with edge width proportional to the extent of the similarity. **C-E** Results from an IPA on the bulk RNA-Seq and proteomics data, always comparing each MC versus all others. Venn diagrams at the top indicate the concordance between RNA and proteomics data in commonly differentially regulated genes/proteins (left) and in pathways found to be significantly enriched in the respective MC (right). Bar plots depict the top scoring pathways found within the RNA-Seq data set, with pathways that are shared with the proteomics data being highlighted in bold italics. Z-Scores are depicted on the x-axis, p-values by the color of the bars.

### Strong enrichment of immune-related pathways in MC ben-1

For a more detailed characterization of the differences between meningioma subclasses, the proteomic and RNA-Seq data sets were subjected to an Ingenuity Pathway Analysis (IPA) based on the differentially expressed genes or proteins in each MC. Interestingly, a strong enrichment in pathways related to the immune system was observed for MC ben-1 compared to the other MCs on both RNA and protein level (Fig. 1C). Tumors of both MC ben-2 and MC mal on the other hand showed a strong signature for MSP-RON signaling in macrophages as well as the PD-1/PD-L1 pathway on both levels (Fig. 1D,E).

Moreover, MC ben-2 tumors were enriched for pathways regulating axon guidance (Reelin signaling pathway, Semaphorin Neuronal Repulsive Signaling Pathway). In addition, the tumor suppressive PTEN signaling pathway displayed increased activity in this MC on mRNA as well as protein level (Fig. 1D).

In MC mal tumors an enrichment of cell cycle related pathways was observed, which is in line with the increased proliferative activity of MC mal tumors (Fig. 1E).

### Single-nuclei RNA-Seq allows characterization of infiltrating cell type composition

In order to investigate whether the strong enrichment for immune response related pathways in MC ben-1 can be traced back to differences in numbers and types of infiltrating cells, an additional 28 meningioma samples from 21 patients were analyzed by single-nuclei RNA-sequencing (10X Single Cell 3’ mRNA Kit v2). Within 26 samples passing QC, all WHO grades and the five most frequent of the six meningioma MCs were represented: seven samples of MC ben-1, three samples of MC ben-2, five samples of MC int-A, three samples of MC int-B and nine samples of MC mal (Suppl. Table 1). The rare ben-3 group, mostly harboring angiomatous meningioma, was not included due to lack of representative samples suitable for isolation.

Individual nuclei, for simplicity in the following denoted as cells, of all samples were clustered in an integrative approach according to similarities in their expression profiles (Fig. 2A). Thereby, multiple clusters of cells with similar expression profiles could be identified, some of which were joint clusters comprising cells from multiple samples while other clusters were unique to individual samples. Using the Human Primary Cell Atlas^11^ as a reference, infiltrating cell types were assigned based on similarity in expression profiles^12^. The obtained cell type annotations were subsequently confirmed through the evaluation of cell typespecific marker gene expression. As a result, one intermixed cluster with cells from multiple samples was classified as macrophages *(MRC1, MS4A7, CD163, LYVE1, STAB1)* as described by us before during neuroinflammation^13–15^. These cells displayed expression profiles characteristic for monocyte-derived macrophages *(CD14, FCGR1A, FCGR3A/B, MRC1)*, although with expression of *P2RY12, SLC2A5*, and *TMEM119* this cell population also exhibited markers typically found in microglia^16^. In addition, one intermixed cluster was identified as lymphocyte cluster comprising T cells (*TRAC*, *TRBC2*, *CD52*, *IL32*) as well as NK cells (*NKG7*, *KLRB1, PRF1, GZMB, GZMA*). Only few B cells (*CD79A, IGHG4, IGLL5*) were present, which clustered together with T cells and NK cells. Furthermore, one endothelial cell cluster (*HSPG2, PLVAP, FLT1, VWF, CD34*), and one mast cell cluster (*CPA3*, *KIT*) were identified (Suppl. Fig. 1A). All remaining clusters were classified as neoplastic cell clusters by expression of meningioma-specific markers, such as *SSTR2*. This classification was further validated with copy number variation (CNV) profiles estimated from the gene expression data (Fig. 2B). Nearly all neoplastic clusters exhibited changes in their CNV profiles, with deletions on chromosome 22q and chromosome 1p being the most common. Only three samples comprised neoplastic cells with no copy number alterations (MNG-17, MNG-18, MNG-20), all of which were of MC ben-2 and displayed flat CNV profiles also in DNA methylation array data (Suppl. Fig. 2A-C). MC ben-2 tumors have previously been shown to differ from the other MCs by virtually flat CNV profiles^8^.

**Figure 2.**
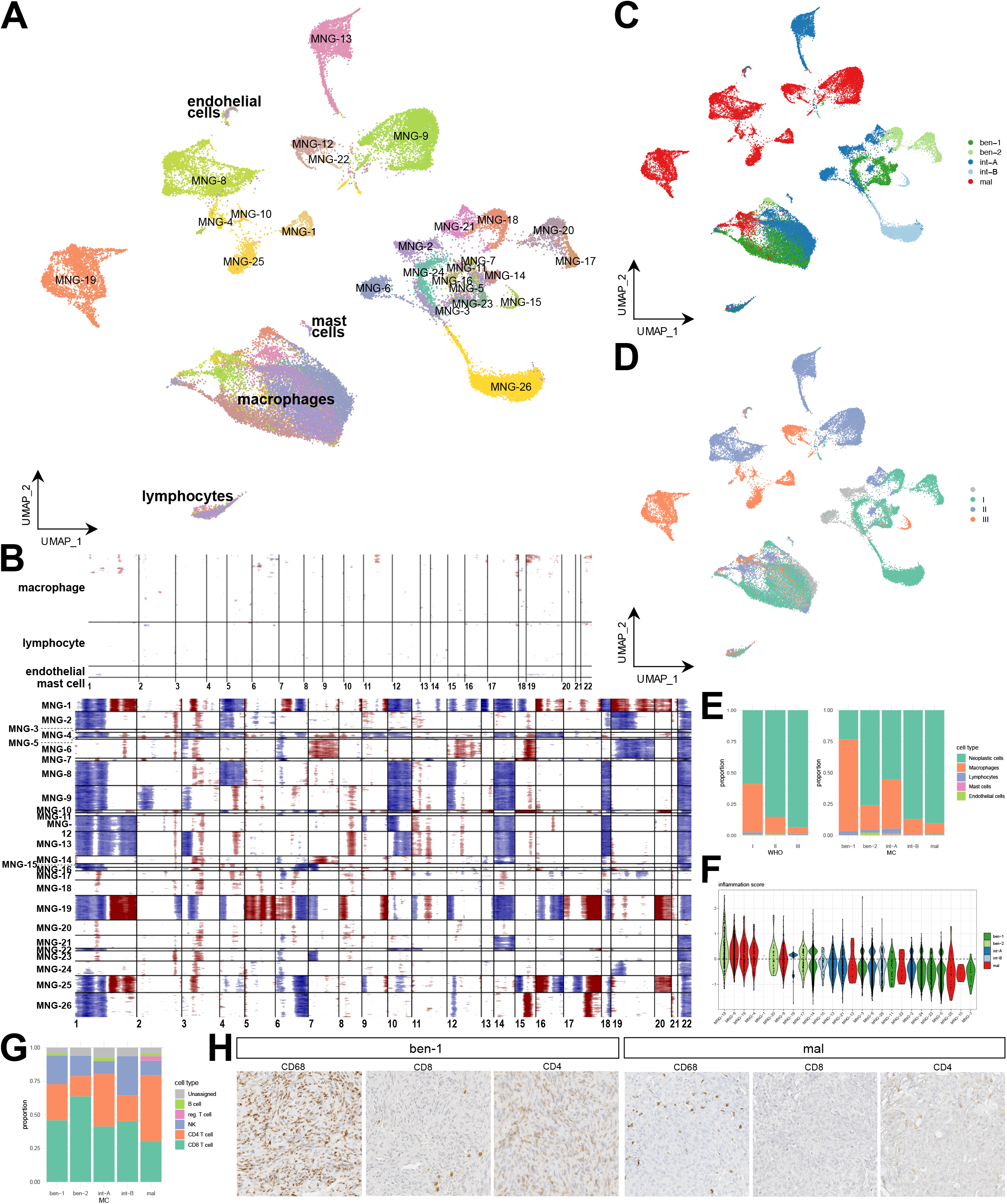
**A** Uniform manifold approximation and projection (UMAP) of the single-nuclei RNA-Seq meningioma dataset comprising 26 samples. Colors indicate the sample of origin for each cell. **B** CNV estimation from the single-nuclei RNA-Seq data for each sample with blue color indicating loss and red color indicating gain at the respective position. At the top, CNV profiles of non-malignant reference cells are shown. **C-D** UMAP from A, with colors representing MCs (C) and WHO grades (D), respectively. **E** Proportions of individual cell types within the single-nuclei RNA-Seq samples assigned to WHO grades (left) and MCs (right). **F** Distribution of inflammation scores of individual macrophages by sample, colors indicating the respective MC. Samples are sorted along the x-axis by decreasing mean inflammation score. **G** Proportions of infiltrating lymphocyte subtypes within the single-nuclei RNA-Seq samples assigned to MCs. **H** Immunohistochemical staining of an MC ben-1 tumor (left) and an MC mal tumor (right) for macrophages (CD68), and CD4+ and CD8+ T-cells, respectively.

The neoplastic cell clusters mostly formed individual clusters unique to samples. Only the WHO grade 1 tumor samples of MC ben-1 clustered more closely together (Fig. 1C,D). The mean distances in the UMAP between samples were smaller for WHO grade 1 and MC ben-1/2 meningiomas compared to the higher-grade tumors (Welch two sample t-test on pairwise centroid distances for samples of WHO 1 vs 3 p-value = 6.06e-4, MC ben-1 vs mal p-value = 7.69e-10, MC ben-2 vs mal p-value = 6.84e-3), indicating a higher inter-sample heterogeneity with increasing malignancy (Suppl. Fig. 1B).

### Infiltrating immune cells differ strongly between MCs

Characterizing the infiltrating immune cells in greater detail, profound differences between WHO grades and MCs in the number and type of infiltrating cells became apparent, in concordance to what was observed from the bulk data. WHO grade 1 tumors displayed slightly larger numbers of infiltrating lymphocytes (Fig. 2E; 1.8 % in WHO grade 1 and 0.9 % in WHO grade 3; Welch two sample t-test on proportions in WHO grade 1 vs grade 3 p-value = 0.0262). The difference in infiltrating macrophages was even clearer, being more abundant in WHO grade 1 compared with higher grade tumors (Fig. 2E; 38.7 %, 12.9 %, and 5.2 % for WHO grade 1, 2, and 3, respectively; Welch two sample t-test on proportions in WHO grade 1 vs grade 3 p-value = 0.0292). When comparing samples by epigenetic group instead of WHO grade, the difference between MC ben-1 and MC mal tumors was yet more pronounced towards higher macrophage infiltration in benign tumors (Fig. 2E; 73.6 % in MC ben-1 and 8.4 % in MC mal; Welch two sample t-test on proportions in MC ben-1 vs MC mal p-value = 0.00520). These results were also validated by immunohistochemistry (51 samples across all MCs, Fig. 2H).

Interestingly, a differential gene expression analysis between the macrophage populations of MC ben-1 and MC mal tumors revealed an upregulation of proinflammatory cytokines (*CCL3, CCL4*) in macrophages infiltrating MC mal tumors (Suppl. Fig. 3A). For further characterization of the macrophage phenotype, macrophages were assigned a proinflammatory score based on the expression of proinflammatory genes (*TNF, IL1B, IL6, IL12A, IL23A, CCL2, CCL8*), and similarly an anti-inflammatory score (*CD163, MSR1, IL10, CD274*). The difference in these scores was termed inflammation score. Macrophages of a proinflammatory phenotype were overrepresented in the MC ben-2, but also the MC mal meningiomas (Welch two sample t-test on proportions in MC ben-1 vs MC mal p-value = 0.0498, and comparing MC ben-1 vs MC ben-2 p-value = 0.0137, respectively; Suppl. Fig. 1E,F). Although the enrichment of proinflammatory macrophages was consistent across samples for MC ben-2, it displayed high variability for MC mal when investigating individual samples (Fig. 2F). While some samples presented a high number of proinflammatory macrophages, others were in the distribution of the inflammation score comparable to the phenotypically more anti-inflammatory MC ben-1 macrophages.

For comparison with the bulk RNA-Seq data, cell populations were estimated from the bulk data through enrichment analysis for gene signatures of the respective cell types. The obtained enrichment scores allow comparisons across samples, although not between cell types within a sample. In line with the single cell data and pathway analysis, enrichment scores for macrophages were highly increased in MC ben-1 (Suppl. Fig. 1C). Applying the inflammation score from the single-nuclei data, which mitigates a bias due to differing total numbers of macrophages by evaluating only the ratio of proinflammatory and antiinflammatory macrophages, a trend of MC ben-2 and MC mal macrophages towards a proinflammatory phenotype was observed, while MC ben-1 macrophages were more inclined towards an anti-inflammatory phenotype (Suppl. Fig. 1D). This was again fully in concordance with the findings from the single-nuclei dataset.

Besides macrophages, lymphocyte types as determined based on hierarchical correlation to a tumor microenvironment reference^17^ also differed between tumor grades and MCs. While an increased number of CD4 T cells was detected in MC mal tumors (Welch two sample t-test on proportions in MC ben-1 and MC mal p-value = 0.0691), the number of CD8 T cells was lower in this class as in lower grade meningiomas (Fig. 2G; Welch two sample t-test on proportions in MC ben-1 and MC mal p-value = 0.00875). NK cells were more abundant in MC mal tumors by trend (Welch two sample t-test on proportions in MC ben-1 and MC mal p-value = 0.0564). Regulatory T cells and B cells were observed only at very low numbers that did not allow for comparisons.

Interestingly, when investigating receptor-ligand interactions between macrophage and lymphocyte populations within each tumor, an increased interaction of macrophages with NK cells through NK cell receptors such as *KLRC2* and *KIR3DL1* was predicted specifically for MC ben-1 tumors (Suppl. Fig. 1G). Furthermore, an IPA comparing macrophage populations between MCs was conducted in a pairwise manner to avoid a bias from the significantly larger numbers of macrophages in MC ben-1 tumors. Thereby, pathways such as Natural Killer Cell Signaling as well as IL-15 Production, an NK cell-activating cytokine, were found to be upregulated in MC ben-1 macrophages (Suppl. Fig. 3B,D). AMPK Signaling, which promotes an anti-inflammatory phenotype in macrophages, was also enriched in macrophages of this MC. Although both MC ben-2 and MC mal macrophages displayed proinflammatory properties, an activation of T and B lymphocytes was observed primarily for MC ben-2 macrophages (Suppl. Fig. 3C,D) while interactions between MC mal macrophages and lymphocytes were less pronounced (Suppl. Fig. 1G,H). The MC mal macrophages instead expressed neutrophil-attracting cytokines such as *IL8* or *IL17F* (Suppl. Fig. 3B).

### CDKN2A expression and FOXM1 activity are enhanced with increasing malignancy

Homozygous deletion of *CDKN2A/B* has recently been introduced as an independent grading criterion for WHO grade 3 meningiomas. Expression levels of *CDKN2A/B* in the single nuclei data set revealed that, surprisingly, across all samples only few cells in the WHO grade 1 and MC ben-1/2 clusters expressed *CDKN2A* (Fig. 3A). In WHO grade 2 and 3, and MC int-A/B and mal, tumors, on the other hand, expression levels of *CDKN2A* were heterogenous between samples: Some (MNG-13) showed high expression whereas in others (MNG-9, MNG-25) the transcript seemed to be absent. All higher-grade tumors with missing *CDKN2A* expression displayed a homozygous deletion of the gene locus as determined on the DNA level (Suppl. Fig. 2D-F). The same finding, namely low *CDKN2A* expression levels in MCs ben-1 and ben-2 and variable expression levels in MC mal, was also observed on RNA as well as protein level in the bulk data (Fig. 3B).

**Figure 3.**
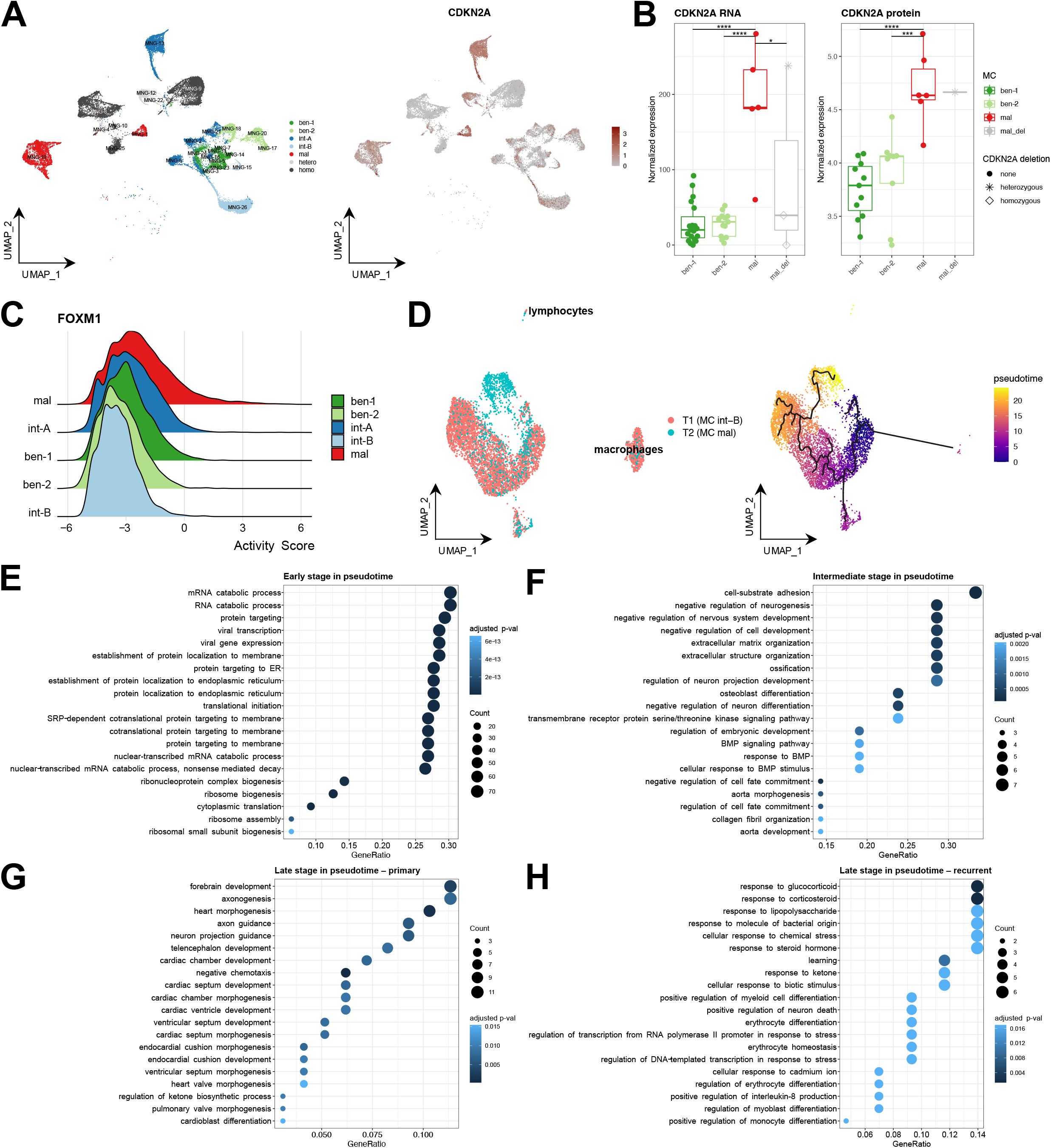
**A** UMAP as in Fig. 2A restricted to the malignant cell population with color indicating MC and status of the CDKN2A gene locus (left) and CDKN2A expression levels (right), where red color indicates elevated expression levels. hetero/homo…MC mal with heterozygous or homozygous CDKN2A deletion, respectively. **B** CDKN2A mRNA (left) and protein (right) expression in meningioma bulk data. *…p-value < 0.05, ***…p-value < 0.001, ****…p-value < 0.0001. **C** Violin plot indicating FOXM1 activity estimated from the expression of its target genes. **D** UMAP of the integrated data from a matched primary and recurrent tumor sample. Colors indicate the sample of origin (left) and the location in pseudotime along an estimated trajectory for the tumor cell population (right). **E-H** Results from a GO enrichment analysis based on the genes expressed early in pseudotime (E), at an intermediate time point (F), or late in pseudotime in the branch specific for the primary (G), and the recurrent tumor (H), respectively.

Another established hallmark of high-grade meningiomas is *FOXM1* network activation. Genes regulated by the *FOXM1* transcription factor have previously been demonstrated to be activated specifically in high grade/MC mal tumors^10,18,19^. From the single-nuclei RNA-Seq data, transcription factor activities were estimated *via* the expression levels of the respective target genes. Thereby, *FOXM1* was indeed found to be inactive in lower grade tumors, while in WHO grade 2 and 3 tumors of MC mal an activation of this transcription factor was observed (Fig. 3C). Interestingly, this activation did not occur homogeneously across all cells of the respective tumor sample, but was rather confined in each case to a small subcluster of cells which also displayed an increased proliferative activity (Suppl. Fig. 3E,F).

### Novel tumor subpopulations emerge in recurrence compared to primary tumor

As recurrences are likely to emerge from a subpopulation of aggressive cells within the primary tumor, neoplastic cell populations were compared between primary and recurrent tumor to identify common and novel cell clusters and their defining features. Therefore, respective samples from a single matched primary and the recurrent tumor pair (primary: WHO grade 1, MC int-B; recurrent: WHO grade 3, MC mal), were integrated by regression of batch effects for a combined analysis (Fig. 3D). Batch effect regression was performed in this case as the biological background can be expected to be more similar in tumors from the same patient. To this end, canonical correlation analysis was performed jointly on both data sets, and mutual nearest neighbors from this representation were used as integration anchors^20^.

A trajectory analysis based on the sequence of changes in gene expression patterns was conducted under exclusion of infiltrating cells. Cells are ordered along this trajectory and annotated with a value in pseudotime based on their position on the trajectory. Thus, the pseudotime reflects the dynamic process which the cell population is undergoing. The cells in the cluster with the least copy number alterations were selected as initial cells and the obtained trajectory revealed a branching point with two alternative end points for cell progression (Figure 3D). Interestingly, one of the end points falls within a cluster mainly made up from cells of the primary tumor, while the second end point is located in a cluster of cells exclusively stemming from the recurrent tumor.

A gene ontology (GO) term enrichment analysis for the marker genes that are differentially expressed and specific for each cell cluster along the trajectory was performed. Cells early in pseudotime displayed a strong enrichment for terms related to translation (Fig. 3E), whereas cells at an intermediate stage in pseudotime exhibited a prominence of terms related to ECM organization and development as well as BMP signaling (Fig. 3F). For the cells late in pseudotime close to the trajectory endpoint within the primary tumor, terms associated with axonogenesis and tissue development were enriched (Fig. 3G), while cells close to the endpoint within the exclusively recurrent tumor cell cluster were enriched for terms connected to stress response and metabolism (Fig. 3H).

### Six tumor cell subgroups with distinct phenotypes can be identified across samples

The findings on the initial Uniform Manifold Approximation and Projection (UMAP) indicate that inter-tumor heterogeneity in meningiomas increases with malignancy. Since the low-grade cases cluster closely together, we next set out to identify traces of common evolution across samples. To find common tumor cell subgroups with similar activities in transcriptomic programs across samples, non-negative matrix factorization (NMF) was applied. This analysis revealed in total six so-called tumor cell meta-clusters with distinct transcriptional signatures, each comprising cells from multiple meningioma samples (Fig. 4A,B). Genes characteristic for each of these meta-clusters were termed signature genes. Based on GO term enrichment analysis for the respective signature genes, different phenotypes were assigned to each meta-cluster. One meta-cluster was mainly comprised of cycling cells, matching well with the cell population that was assigned to the G2M/S phase previously (Suppl. Fig. 3F), whereas another meta-cluster displayed elevated expression levels of genes connected to translational activity. The remaining meta-clusters were enriched for GO terms related to synapse assembly, myelination, extra-cellular matrix (ECM) organization, or angiogenesis, respectively (Suppl. Fig. 4A). Intriguingly, cells of the different MCs were not evenly distributed across the meta-clusters: The majority of cells in the meta-clusters of cell cycle activation, ECM organizing, and angiogenic tumor cells stemmed from MC mal meningiomas, whereas the synaptogenesis related meta-cluster was dominated by cells from benign and intermediate meningiomas (Fig. 4C). The translational and myelination related meta-clusters displayed no clear difference in their distribution across MCs. The synaptogenesis and myelination related meta-clusters were the only meta-clusters present across all samples, however to smaller extent in the MC mal tumors. A gene set enrichment analysis based on the scoring of each gene in the respective meta-cluster indicated an increased expression of genes related to VEGFA as well as TGFβ signaling in the angiogenesis meta-cluster (Suppl. Fig. 4C), while the synapse assembly-related meta-cluster was enriched for the PI3K/AKT signaling pathway, but also the MAPK signaling pathway (Suppl. Fig. 4B).

**Figure 4.**
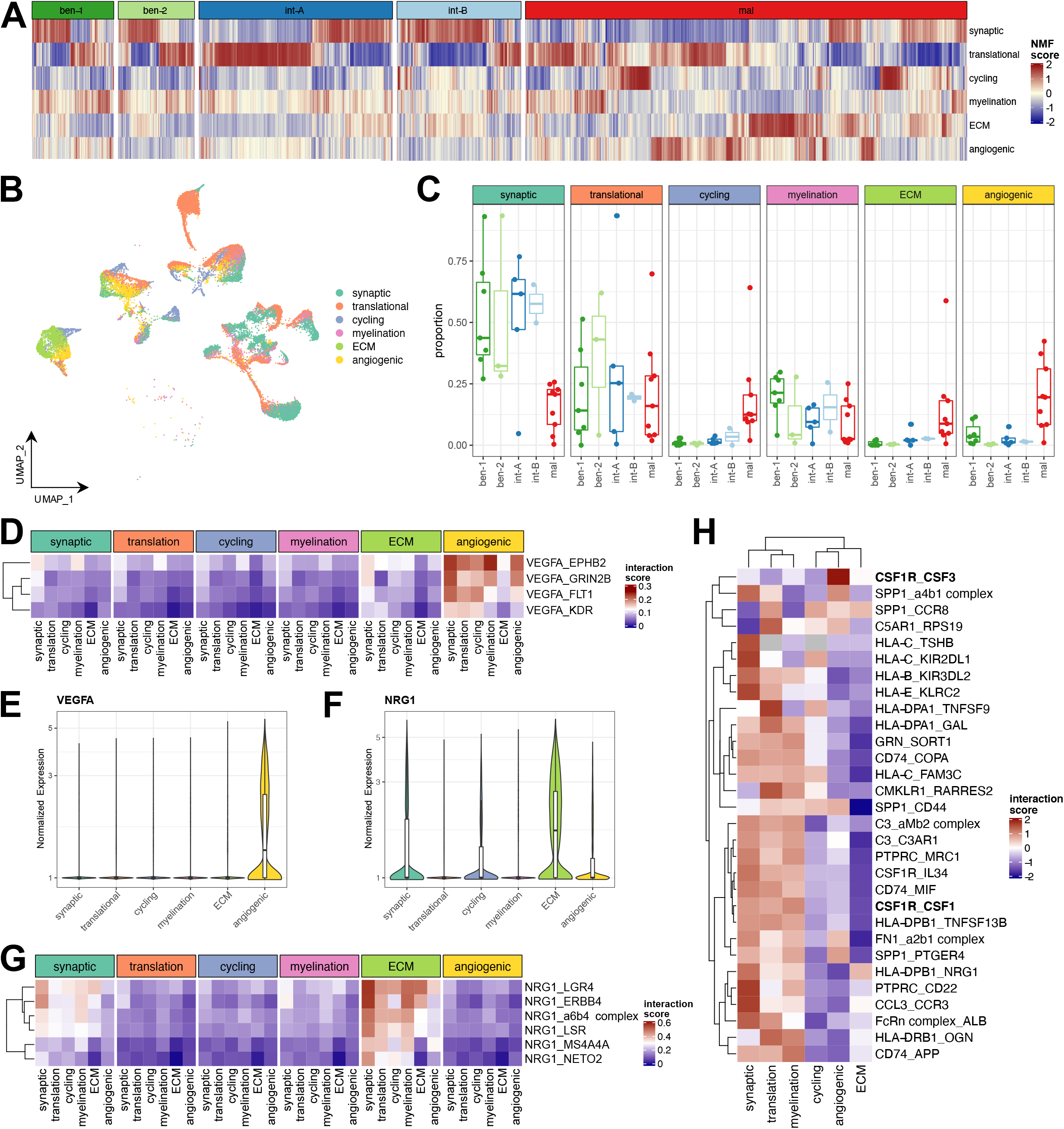
**A** Heatmap indicating the enrichment for each of the six meta-signatures per cell (column) sorted by MC. Red color indicates high score for the respective NMF meta-cluster, blue indicates low score. **B** UMAP from Fig. 2A, restricted to the malignant cell population, with colors representing the assignment of each cell to one of six identified meta-signatures. **C** Proportion of cells per sample assigned to each of the meta-clusters sorted by MC. **D** Heatmap of interaction scores between cells of the NMF meta-clusters involving *VEGFA*. Top annotation indicates source of the interaction (expressing the ligand), bottom annotation indicates target (expressing the receptor). **E,F** Violin plots indicating *VEGFA* (E) and *NRG1* (F) expression per NMF metacluster, respectively. **G** Heatmap of interaction scores between cells of the NMF meta-clusters involving *NRG1*. Top annotation indicates source of the interaction (expressing the ligand), bottom annotation indicates target (expressing the receptor). **H** Heatmap of interaction scores for cells in each NMF meta-cluster with the infiltrating macrophage population for the 30 ligandreceptor pairs with highest variability in interaction scores across meta-clusters.

Through investigation of the receptor-ligand interactions between meta-clusters, inferred from matching expression patterns of ligand and receptor in a pair of clusters^21^, the metacluster of the angiogenic phenotype was found to act through activation of *VEGFA* receptors on all other meta-clusters, except the ECM organization cluster (Fig. 4D). This was in concordance with the elevated *VEGFA* expression levels observed for this cluster (Fig. 4E). Moreover, the ligand-receptor interaction analysis revealed a strong signaling signature and elevated expression levels of *NRG1* in the ECM organizing meta-cluster (Fig. 4F,G), mainly consisting of cells from meningiomas of MC mal. *NRG1* signaling through *ERBB4* can activate the MAPK as well as the AKT/PI3K signaling cascade. *NRG1* has previously been shown to be secreted at high levels in meningiomas upon loss of *NF2*^22^. This is in line with the finding that virtually no cells from MC ben-2 meningiomas, which do not have this deletion, were represented in this meta-cluster.

Similarly, interactions of the tumor meta-clusters with the immune cell populations, specifically the infiltrating macrophages, were investigated in greater detail. Types and degree of interaction with the macrophages differed strongly between meta-clusters. Strongest interactions were observed for the meta-clusters with synaptic, translational, and myelinating signatures enriched in benign and intermediate MCs (Fig. 4H). Interestingly, cells in these meta-clusters seemed to stimulate macrophages via their *CSF1R* by secreting *CSF1*, while cells in the angiogenic meta-cluster secreted *CSF3* acting on the same receptor (Fig. 4H).

### Phosphoproteomic analyses reveal MAPK/ERK pathway activity restricted to benign meningiomas

To investigate activated pathways in greater detail, phosphoproteomics data was generated for ten samples representing WHO grades 1 and 3, and MCs ben-1, ben-2, and mal. Repeating the SNF analysis for the eight samples with MCs ben-1/2 and mal with complete data, here also including the phosphoproteomics layer in addition to proteomics and RNA level, illustrated a similar pattern as above with MCs clustering separately and MC ben-2 and mal more closely related (Suppl. Fig. 5A).

A subsequent kinase perturbation analysis revealed elevated activities for several kinases of the MAPK family (*MAPK1*/ERK2, *MAPK3*/ERK1, *MAPK14*/p38α) in MC ben-1 as well as ben-2, clearly indicating an activation of the MAPK/ERK pathway in these meningiomas (Suppl. Fig. 5B). This is in line with previous findings, where low grade meningioma displayed an increased activation of the MAPK pathway compared to higher grade meningiomas^23^.

In addition, *AKT1* displayed activity exclusively in MC ben-2 tumors (Suppl. Fig. 5B), reflecting frequent *AKT1/PIK3CA* mutations in this MC, and indicating an activated AKT/PI3K signaling pathway.

MC mal tumors on the other hand showed elevated activity levels for cell cycle regulating kinases (*CDK2, CDK7*), consistent with their increased proliferative activity (Suppl. Fig. 5B).

## DISCUSSION

The landscape of meningioma can be dissected into the continuum of *NF2* mutant cases from high to low grade on the one, and the non-*NF2* cases, mostly low grade, on the other hand^3,4,7^. Several stratification approaches into clinically meaningful subgroups have been proposed based on these molecular markers, including DNA methylation-based MCs^8^. Here, we demonstrate that these MCs delineate biological groups that are consistently recapitulated across epigenomic, transcriptional and proteomic levels. These findings support the role of MCs as robust subtyping approach in meningioma classification.

Within the MCs, the differences in proportions of infiltrating lymphocytes and even more so of infiltrating monocyte-derived macrophages is intriguing. As this was also validated by computational decomposition of bulk RNA-Seq data as well as immunohistochemistry, a technical artifact during nuclei extraction can be excluded. The highly MC-specific distribution of lymphocytes, along with an activation of the immune checkpoint axis in some MCs (MC ben-2 and MC mal), may provide the rationale for intensified studies on immunotherapy in meningioma.

The expression profile of the monocyte-derived macrophage cell population, on the other hand, does not fully conform with macrophages, but in addition exhibits microglial markers. Meningiomas are non-parenchymal tumors of the CNS, and none of the meningiomas included here presented with brain invasion. Therefore, an infiltration with actual microglia seems unlikely based on current knowledge, even when considering concepts of CNS/meningeal lymphatics^24–27^. Hence, the expression profile may suggest the presence of a specific macrophage subset. CX3CR1+ resident macrophages are present in the dura and arachnoidea mater and specifically also found in the choroid plexus^28^, another location in which *NF2* mutant meningiomas are frequent^29^. The enrichment of *NF2* mutant meningioma in the lateral ventricles is consequently in line with the association of *NF2* mutations/*NF2* mutant subtype MC ben-1 and CX3CR1+ macrophages, while a non-convexity localization is typical for non-*NF2* mutant meningiomas.

However, not only abundance, but also activation of cells within the tumor microenvironment differed between MCs. Macrophages of MC ben-1 tumors are capable of activating NK cells, thus possibly aiding in preventing rapid tumor growth as seen in high grade tumors. By contrast, macrophages in MC mal meningiomas preferentially attracted neutrophils, probably due to necrotic areas occurring in high grade tumors. Given these differences in numbers and activation patterns of lymphocytes and macrophages between low- and high-grade tumors, infiltrating immune cells might play an important role in determining the malignancy of a meningioma.

In search of further interdependence of tumor growth and myeloid cell infiltration, we found that especially the tumor subpopulations dominated by cells from low-grade tumors stimulated macrophage production and differentiation via *CSF1*. As this gene is located on chromosome 1p, a gene locus frequently deleted in high-grade meningioma, absence of *CSF1* could be one factor in MC mal tumors contributing to the decreasing numbers of infiltrating macrophages with increasing malignancy. Accordingly, in MC int-A, for which cases with and without chromosome 1p deletion were available, a decrease in *CSF1* expression was observed in the 1p deleted cases. Controversially, elevated *CSF1* expression levels have been associated with an unfavorable outcome in various tumor diseases^30–33^. Along these lines, a recent study found elevated levels of *CSF1* in the plasma of meningioma patients, a correlation of *CSF1* expression and an increased number of infiltrating macrophages of an anti-inflammatory phenotype, and a beneficial effect of *CSF1/CSF1R* targeting antibody treatment in a meningioma mouse model^34^.

Analyzing single nuclei from meningioma cells across the different WHO grades and MCs, we were able to define several tumor subpopulations with distinct phenotypes, with varying abundance depending on tumor grade. Present in all samples but significantly more prevalent in lower-grade tumors, are cells with a synaptogenesis related phenotype and activity in the MAPK as well as AKT/PI3K signaling pathway. A similar phenotype was observed in bulk data on transcriptional and translational level for MC ben-2 tumors. A second subpopulation with an ECM organizing phenotype was predicted to interact with these cells via secretion of *NRG1*, which may activate both MAPK and AKT/PI3K signaling through *ERBB4* receptors. This subpopulation was mainly present in high-grade tumors and absent in MC ben-2 tumors without *NF2* loss, which has been shown to induce elevated expression of *NRG1*^22^. A malignant cell population of similar phenotype was found at an intermediate stage of a trajectory for a matched primary and recurrent tumor pair, before branching of and giving rise to a cell population dominated by stress response exclusive to the recurrent tumor. This stress response, and signals of DNA repair, are in line with increased proliferation, but the emergence of hypoxia also has a correlate in the morphologically detected necrosis in these tumors. In turn, the proliferative stress in these cells may be the cause for the accumulation of chromosomal alterations and genome-wide instability in malignant meningiomas^8,10^. Similarly, an angiogenic tumor subpopulation with cells under hypoxic stress with TGFβ signaling activity was identified and expectedly enriched in MC mal samples, further substantiating previous observations of TGFβ activation in malignant meningioma and supporting the existing efforts of TGFβ blockade in meningiomas^35–37^.

Previous studies have proposed *FOXM1* network activation as an underlying mechanism of malignant transformation in meningioma. In support of this mechanism, our data indicate the development of a *FOXM1*-activated subset of cells within lower grade meningiomas, giving rise to a high-grade tumor. Hence, these findings emphasize the role of *FOXM1* activation as an early marker and potential treatment target in high-grade meningioma. Surprisingly, the expression of the gene product of *CDKN2A/B*, p16, was not uniformly altered in high grade compared to low grade cases. While the homozygous deletion of *CDKN2A/B* is a novel marker for anaplasia in the upcoming WHO classification of brain tumors, its expression increased from low-grade to high-grade cases, consistently on transcriptomic and proteomic levels, except for cases with homozygous deletion. This may indicate a cell-intrinsic barrier to further malignant transformation, while exposing the highly translated, “open” *CDKN2A/B* locus to rearrangements and deletion events. Clinically, this concept of a further segregation between high-grade *CDKN2A/B* intact and homozygously deleted cases is in line with the even more unfavorable outcome of the latter. From a diagnostic perspective, this observation may also explain why p16 immunohistochemistry has so far not proven useful in grading of meningioma.

Collectively, we identified molecular subgroups of meningioma that are robust throughout several molecular layers from epigenetic regulation to translation, emphasizing their relevance for biological and clinically meaningful classification, and suggest subtype-specific pathway and immune activation as potential novel treatment targets.

## ACKNOWLEDGEMENTS

This study was supported by the Geman Cancer Aid (70112956) and Else Kröner-Fresenius Foundation (EKFS, 2015_A60 and 2017_EKES.24). CB is a scholar of the DKFZ International Graduate School. Philipp Sievers is a fellow of the Hertie Network of Excellence in Clinical Neuroscience. We thank Laura Dörner, Lea Hofmann and Moritz Schalles for skillful technical assistance.

## METHODS

### Tissue selection

Tissues were selected from the archives of the Dept. of Neuropathology and Neurosurgery Heidelberg, Mannheim, Paris and Zurich based on tissue availability and in order to cover all WHO grade and MCs (2018-614N-MA, 005/2003). Especially for frozen tissue, quality was determined by frozen sections.

### Bulk RNA-Sequencing

RNA-Seq was performed as previously described38. In short, library preparation was performed with the TruSeq RNA Library Prep for Enrichment kit (Illumina) and paired-end reads were sequenced on a NextSeq 500 instrument (Illumina). After adapter trimming, reads were aligned to the human genome (GRCh37) with the STAR aligner39 and counted using RSEM^40^. Differential expression analysis was performed in R v.4.0.0^41^ using the DESeq2 package^42^. P-values for differentially expressed genes were adjusted for multiple testing with the Benjamini-Hochberg procedure.

### Proteomics and Phosphoproteomics

FFPE tissue was deparaffinized and prepared as described recently^43^. To generate reference libraries for data-independent analyses, one μg of each sample was pooled and separated into 16 fractions using high pH reversed-phased pre-fractionation^44^. Phosphorylated peptides were enriched on an AssayMAP Bravo automation platform (Agilent Technologies) using 5 μL Fe(III)-NTA cartridges (Agilent, G5496-60085) according to the instructions of the manufacturer. Subsequently, liquid chromatography (LC) - mass spectrometry (MS) measurements were conducted either on a QExactiveTM HFX Orbitrap (Thermo Fisher Scientific) or a timsTOFPro (Bruker Daltonics) instrument coupled to an EASY-nLC 1200 ultrahigh-pressure system (Thermo Fisher Scientific). MS data were acquired using data-independent (DIA) or data-dependent (DDA) modes employing acquisition parameters as defined earlier^43,45^. MS data were processed in a hybrid approach combining the SprectromineTM (version 1.0.21621.11.1692, Biognosys) and SpectronautTM (version 13.1.190621.43655, Biognosys) software. Phosphoproteomic data were analyzed in MaxQuant (v 1.6.17.0). All searches were accomplished using the human Uniprot reference databases UP000005640_9606 and UP000005640_9606_additional. For phosphoproteomic data, Phospho (STY) were additionally specified as sites of variable modification.

### DNA methylation analysis

All methylation data were generated using the Illumina MethylationEPIC (850k) array platform according to the manufacturer’s instructions (Illumina). Sample preparation was performed as previously described^46^. DNA methylation status of 10,000 CpG sites was analyzed on the current version v12b4 of the Classifier (https://www.molecularneuropathology.org/mnp).

### Similarity network fusion for RNA-Seq and proteomics

Similarity Network Fusion (SNF) was applied to RNA-Seq, proteomics, and, in a second set, phosphoproteomics data if available^47^. No prior feature selection was conducted, but instead Euclidean distances between samples were calculated from the full feature matrices. From these, first affinities were calculated using the R package SNFtool^48^ with the number of nearest neighbors set to ten (and three for the smaller sample set including the proteomic data) and the hyperparameter alpha set to 0.5 and affinity matrices were subsequently fused into one consensus network with the number of iterations for the diffusion process set to ten. In the resulting consensus network weak similarities are removed while weights of connections supported by similarities in several data modalities are increased.

### Ingenuity pathway analysis (IPA)

For pathway analyses, differentially expressed genes and proteins, respectively, were calculated in a pairwise fashion between MCs. Genes and proteins with an adjusted p-value < 0.05 were considered significantly differentially expressed. From the RNA-Seq data, only genes with an absolute log fold change > 1 were included in the pathway analysis. IPA (QIAGEN Inc., https://www.qiagenbioinformatics.com/products/ingenuity-pathway-analysis) was conducted based on fold changes and pathways with a p-value < 0.05 were considered significant.

### Single Nuclei Extraction

Single nuclei suspensions were obtained from frozen tumor tissues as described by Ernst et al.^49^ with some modifications. Briefly, the tumor content was assessed on a Hematoxylin and eosin stain and a fragment of 30-40 mg tissue was cut in the same orientation. The tissue was mechanically lysed with a scalpel in a total volume of 5 ml lysis buffer (0.32 M sucrose [Sigma-Aldrich 84097], 5 mM calcium dichloride [Sigma-Aldrich 21115], 3 mM magnesium acetate [Sigma-Aldrich 63052], 2.0 mM EDTA [Invitrogen 15575-038], 0.5 mM EGTA [Alfa Aesar J61721], 10 mM Tris-HCl, pH 8.0 [Invitrogen AM98556], 1 mM DTT [Sigma-Aldrich 10197777001] and 0.1% Triton X-100 [Sigma-Aldrich 93443]). Next, the suspension was transferred into a glass douncer (Sigma-Aldrich D9063) and further lysed by douncing eight strokes each with pestle A and B. The lysate was directly filtered through a 100 μm filter (Greiner Bio-One 542000) followed by a 40 μm filter (Greiner Bio-One 542040) into a precooled and coated Falcon tube. After spinning (500g, 5 min at 4°) and washing for a maximum of three times (wash buffer: lysis buffer w/o Triton X-100 and DTT), the nuclear pellet was resuspended in a volume of 1 ml storage buffer (0.43 M sucrose [Sigma-Aldrich 84097], 70 mM potassium chloride [ThermoFischer Scientific AM9640G], 2 mM magnesium dichloride [ThermoFischer Scientific AM95306], 10 mM Tris-HCl, pH 7.2 [Sigma-Aldrich T2069] and 5 mM EGTA [Alfa Aesar J61721]). Subsequently, the nuclei suspension was mixed by using a 200 μl pipette to minimize clump formation. The final nuclei number was quantified on a bright-field automated cell counter after staining an aliquot with trypan blue at 1:1 (Luna-BF, Logos Biosystems).

### Single-nuclei RNA-Seq Library Preparation & Sequencing

Single-nuclei RNA-seq libraries were prepared according to the Chromium Next GEM Single Cell 3’ Reagent Kits v2 User Guide (10x Genomics). After counting, nuclei concentrations were adjusted to the desired capture number based on the number of available nuclei. A slightly higher number of nuclei were used to compensate losses in subsequent steps. To minimize potential multiplets, we typically aimed to capture around 5,000 nuclei per sample. For cDNA amplification a total number of 14 cycles was set. QC of the cDNA as well as the final sequencing libraries were performed on the TapeStation 4200 platforms. Concentrations were determined on a Fluorometer (Qubit 4 Fluorometer, Thermo Fisher Scientific) with the Invitrogen Qubit DNA HS Assay Kit (Thermo Fisher Scientific, Q32854).

The final single indexed sequencing libraries were loaded on a NovaSeq 6000 (Illumina) using 100 cycles kits and the following read lengths: 28 bp Read 1, 8 bp i7 index, 0 bp i5 index and 94 bp Read 2. The estimated saturation of detected nuclei was at approximately 20,000 reads per nucleus.

### Single-nuclei RNA-Seq analysis

Single-nuclei RNA-Seq data was aligned to the genome and quality controlled with Cell Ranger v.3.0.1 (10X Genomics). Cells with less than 200 detected features or a median absolute deviation of more than three times in their detected features were excluded from the analysis. Also, cells for which the percentage of mitochondrial genes was higher than three times the median value within a sample were filtered out. All analyses were performed in R v.4.0.0^41^ with the Seurat package v.4.0.0^50^ unless otherwise specified.

The data was normalized and scaled and the number of counts per cell was regressed out. Cell type annotation was performed using SingleR^12^ with the Human Primary Cell Atlas^11^ as reference, and in addition based on the expression of cell type specific marker genes. Tumor cells were identified based on CNV profiles. The CNV profiles were estimated using the InferCNV R package^51^. To this end, within each sample expression profiles of five cells within the same cluster were averaged, and for the thereby obtained meta-cell the CNV status was determined by applying a sliding window across 200 neighboring genes within each chromosome and median filtering for smoothing.

Cell-cell interactions were calculated for each sample individually with CellPhoneDB^21^ based on the expression levels of a receptor and its respective ligand in two defined cell populations.

### Trajectory analysis of matched primary and recurrent single-nuclei RNA-Seq sample

Regression of batch effects was performed for the matched primary and recurrent tumor pair in Seurat^20^. Therefore, both data sets were submitted jointly to a canonical correlation analysis. This representation was used to determine nearest neighbors, which served as integration anchors.

Trajectory analysis was performed using the Monocle 3 package^52^. The trajectory represents a sequence of changes in gene expression, calculated on the UMAP reduction of the data set. Cells with less alterations in their CNV profile were subsequently chosen as starting cells to order the cells along a pseudo time with the starting cells at pseudo time point zero and the remaining cells ordered based on the difference in expression patterns. Genes specific for a certain stage in pseudo time were determined by comparing gene expression in tumor cells at this time point to the expression at all other time points.

### Identification of tumor subpopulations with common transcriptional programs

To identify shared signatures in the expression profiles of tumor cells across samples, NMF was applied as described. To this end, expression data for the tumor population of each sample was centered and scaled before performing NMF with a factorization rank of three for each sample individually^53^. For all resulting factors, the top 30 genes with highest NMF scores were selected as the gene signature specific for that factor and signatures were scored in each cell. Scoring was performed by first determining those 100 genes with highest difference in mean expression across the whole data set to the mean expression of the respective gene and subsequently calculating the difference in expression of the respective gene to the mean expression of those selected 100 genes in each cell. Scores for all genes in a signature were averaged to obtain scores for the gene signature of all factors. These signatures were then hierarchically clustered based on their scores per cell. This revealed six correlated signature sets. Cells were assigned to one of those six so-called meta-programs on the basis of the highest average score in the respective cell. Genes specific for the metaprograms were determined as the 50 genes with highest average score in the cells belonging to that meta-program.

### Immunohistochemistry

Immunohistochemistry was performed on a BenchMark XT immunostainer and on a BenchMark Ultra immunostainer (Ventana Medical Systems). Therefore, 0,5-μm-thick formalin-fixed, paraffin-embedded (FFPE) tissue sections were mounted on SuperFrost Plus Adhesive slides (Thermo Scientific) followed by drying at 75 °C for 10 min.

Dilutions and antibody details are provided in Suppl. Table 2.

Slides were scanned on an Aperio AT2 Slide Scanner (Biosystems Switzerland AG) and analyzed using Aperio ImageScope software (v11.0.2.725, Aperio Technologies).

### Estimation of cell type enrichment scores from bulk RNA-Seq data

Cell type proportions of infiltrating immune and stroma cells were estimated through an enrichment analysis for cell type-specific expression profiles with xCell54. The enrichment of the gene signature for a respective cell type in a bulk RNA-Seq dataset was compared across samples and resulting enrichment scores were transformed to a linear scale. Scores were averaged across samples within the same MC.

---

**Supplementary Figure 1. A** UMAP as in Fig. 2A with color indicating the cell type specific marker gene expression for macrophages, T cells, NK cells, B cells, endothelial cells, and mast cells. Blue color indicates low score for expression, yellow color indicates high score. **B** Mean Euclidean distances between cluster centroids of the tumor cell population from the UMAP for each pair of samples within the same WHO grade (left) and MC (right), respectively. **C** Enrichment of infiltrating cell types in MCs ben-1, ben-2, and mal as estimated from bulk RNA-Seq data with xCell. **D** Inflammation score estimated from the bulk RNA-Seq data based on the difference in mean expression of proinflammatory genes and mean expression of antiinflammatory genes. **E** UMAP of the macrophage subpopulation after regression for batch effects between samples. Colors indicate tumor of origin. **F** UMAPs as in E split by MC, with colors indicating the expression of proinflammatory marker genes. **G,H** Heatmap of interaction scores between macrophage and lymphocyte populations within each tumor averaged by MC with macrophages as source of the interaction (expressing the ligand, G) or as target (expressing the receptor, H). ***…p-value < 0.001, ****…p-value < 0.0001.

**Supplementary Figure 2.** CNV profiles for three tumors of the single-nuclei RNA-Seq dataset estimated from DNA methylation array. **A-C** Flat CNV profiles of MC ben-2 tumors. **D-E** Status of the CDKN2A gene locus. In two samples (D,F) the CDKN2A locus is deleted, whereas it is intact in the remaining sample (E).

**Supplementary Figure 3. A** Differentially expressed genes between the macrophage populations of MC ben-1 and MC mal tumors. **B-D** Ingenuity Pathway Analysis (IPA) comparing macrophage populations between MCs ben-1, ben-2, and mal in a pairwise fashion. **E** UMAP as in Fig. 2A restricted to the malignant cell population with color indicating FOXM1 activity as estimated from target gene expression. Red indicates high score for activity, blue indicates low score. **F** UMAP as in Fig. 2A with color indicating the phase in the cell cycle for the respective cell.

**Supplementary Figure 4. A** Results from a GO enrichment analysis based on the signature genes identified for each of the six NMF meta-clusters. **B,C** Gene set enrichment analysis based on the gene scores from the NMF analysis for the synapse assembly (B) and angiogenesis related meta-cluster (C).

**Supplementary Figure 5. A** SNF graph deducted from bulk RNA-Seq, proteomics, and phosphoproteomics data. Each node represents a sample with colors indicating the respective MC. Edges between nodes indicate similarity between the sample with edge width proportional to the extent of the similarity. **B** Results from a kinase perturbation analysis, calculated based on the phosphoproteomics data as comparison of each MC versus all remaining samples. Red indicates high predicted activity of the respective kinase, blue indicates low activity. Asterisks mark significant kinases. Only kinases with at least three identified targets were included.

## SUPPLEMENT

## Abbreviations

CNS: Central nervous system
CNV: Copy number variations
ECM: Extracellular matrix
IPA: Ingenuity pathway analysis
GO: Gene ontology
MC: Methylation class
NMF: Non-negative matrix factorization
SNF: Similarity network fusion
RNA-Seq: RNA-Sequencing
UMAP: Uniform Manifold Approximation and Projection
WHO: World Health Organization

---

**Suppl. Table 1.**
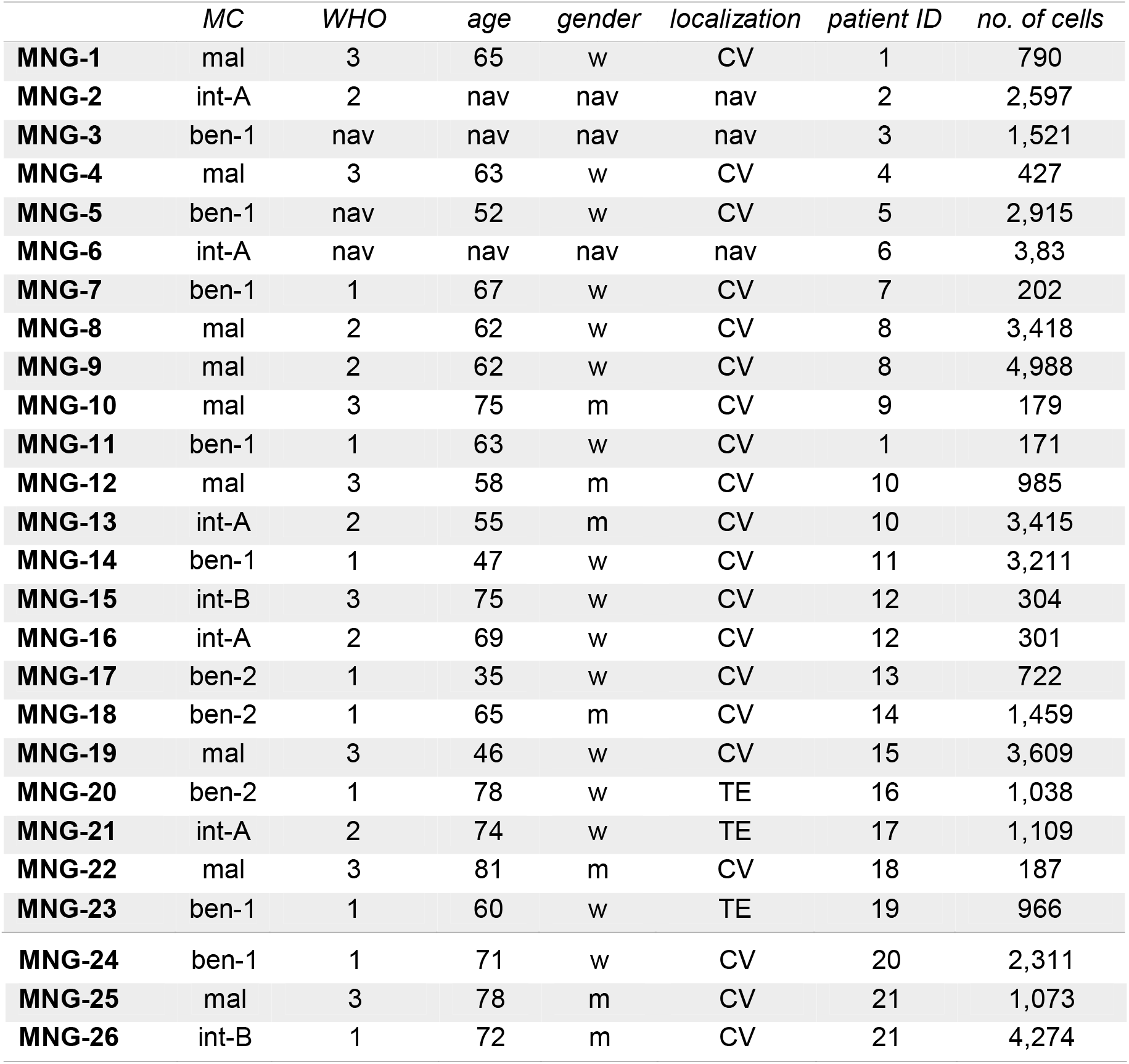
Single-nuclei RNA-Seq sample overview

|  | *MC* | *WHO* | *age* | *gender* | *localization* | *patient ID* | *no. of cells* |
| --- | --- | --- | --- | --- | --- | --- | --- |
| **MNG-1** | mal | 3 | 65 | w | CV | 1 | 790 |
| **MNG-2** | int-A | 2 | nav | nav | nav | 2 | 2,597 |
| **MNG-3** | ben-1 | nav | nav | nav | nav | 3 | 1,521 |
| **MNG-4** | mal | 3 | 63 | w | CV | 4 | 427 |
| **MNG-5** | ben-1 | nav | 52 | w | CV | 5 | 2,915 |
| **MNG-6** | int-A | nav | nav | nav | nav | 6 | 3,83 |
| **MNG-7** | ben-1 | 1 | 67 | w | CV | 7 | 202 |
| **MNG-8** | mal | 2 | 62 | w | CV | 8 | 3,418 |
| **MNG-9** | mal | 2 | 62 | w | CV | 8 | 4,988 |
| **MNG-10** | mal | 3 | 75 | m | CV | 9 | 179 |
| **MNG-11** | ben-1 | 1 | 63 | w | CV | 1 | 171 |
| **MNG-12** | mal | 3 | 58 | m | CV | 10 | 985 |
| **MNG-13** | int-A | 2 | 55 | m | CV | 10 | 3,415 |
| **MNG-14** | ben-1 | 1 | 47 | w | CV | 11 | 3,211 |
| **MNG-15** | int-B | 3 | 75 | w | CV | 12 | 304 |
| **MNG-16** | int-A | 2 | 69 | w | CV | 12 | 301 |
| **MNG-17** | ben-2 | 1 | 35 | w | CV | 13 | 722 |
| **MNG-18** | ben-2 | 1 | 65 | m | CV | 14 | 1,459 |
| **MNG-19** | mal | 3 | 46 | w | CV | 15 | 3,609 |
| **MNG-20** | ben-2 | 1 | 78 | w | TE | 16 | 1,038 |
| **MNG-21** | int-A | 2 | 74 | w | TE | 17 | 1,109 |
| **MNG-22** | mal | 3 | 81 | m | CV | 18 | 187 |
| **MNG-23** | ben-1 | 1 | 60 | w | TE | 19 | 966 |
| **MNG-24** | ben-1 | 1 | 71 | w | CV | 20 | 2,311 |
| **MNG-25** | mal | 3 | 78 | m | CV | 21 | 1,073 |
| **MNG-26** | int-B | 1 | 72 | m | CV | 21 | 4,274 |

**Suppl. Table 2.** Antibodies and dilutions used for immunohistochemical stainings.

| *Antibody* | *Manufacturer* | *Clone* | *Catalogue No.* | *Pretreatment* | *Antibody dilution* | *Antibody incubation* |
| --- | --- | --- | --- | --- | --- | --- |
| CD4 | Cell Marque | SP35 | 104R-16 | CC2 (pH=6,0), 56min | 1:50 | 32min |
| CD8 | Dako | C8/144B | M7103 | CC1 (pH=8,4), 52min | 1:50 | 32min |
| CD68 | Dako | PG-M1 | M0876 | CC1 (pH=8,4), 36min | 1:20 | 32min |

