## Supplementary figures and images for "Integrated phospho-proteogenomic and single-cell transcriptomic analysis of meningiomas establishes robust subtyping and reveals subtype-specific immune invasion"

### Suppl. Fig. 1

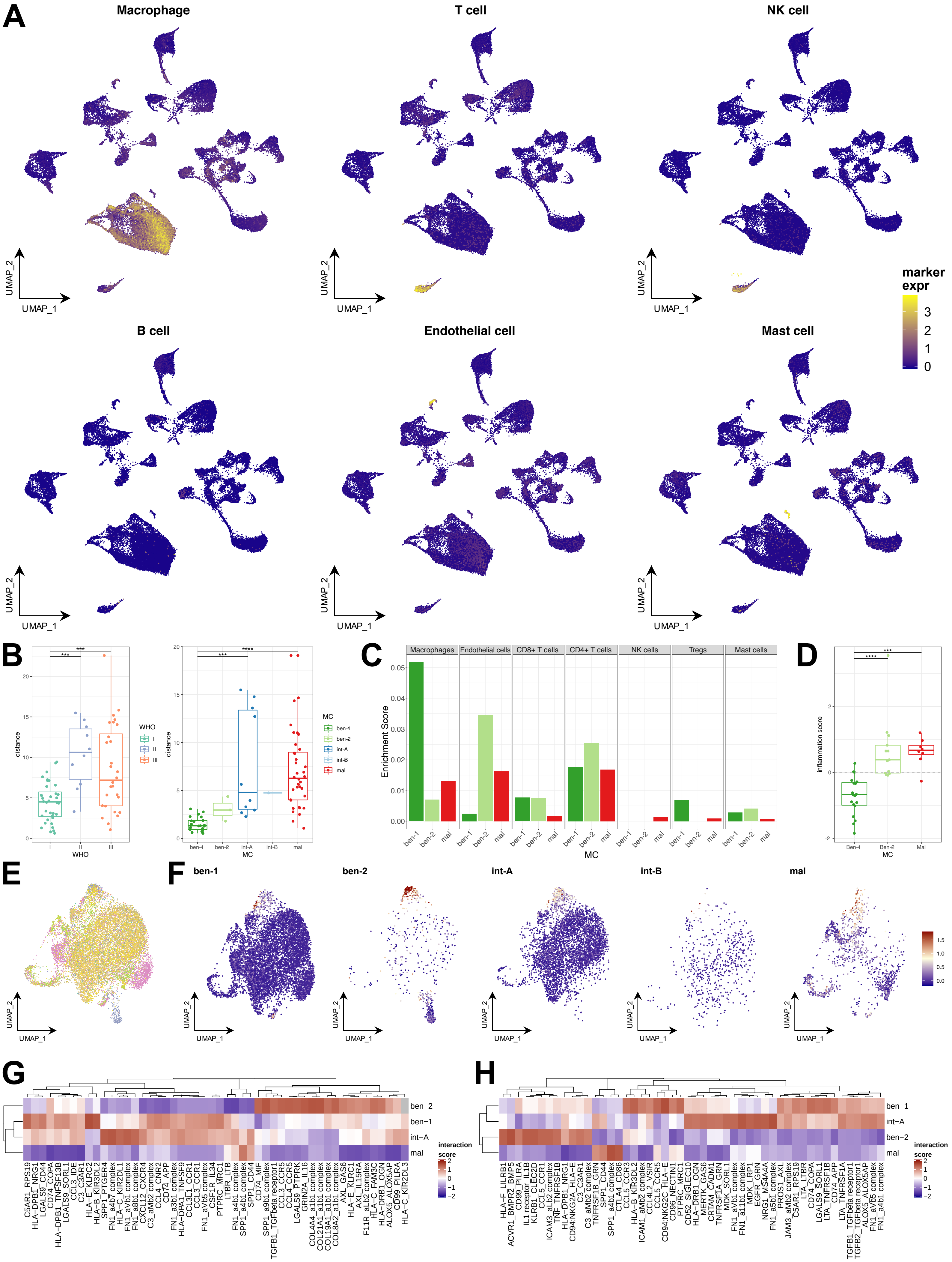

### Suppl. Fig. 2

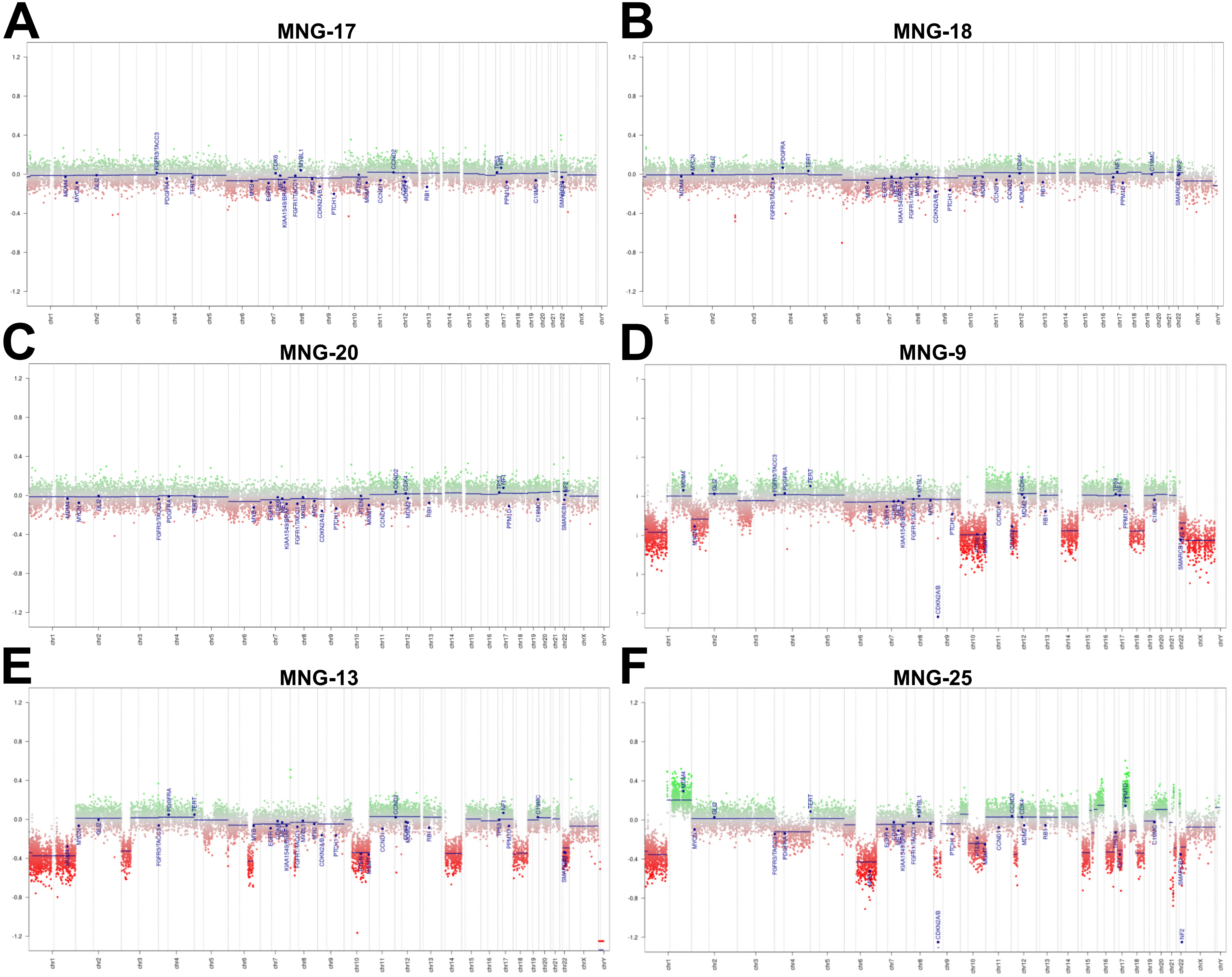

### Suppl. Fig. 3

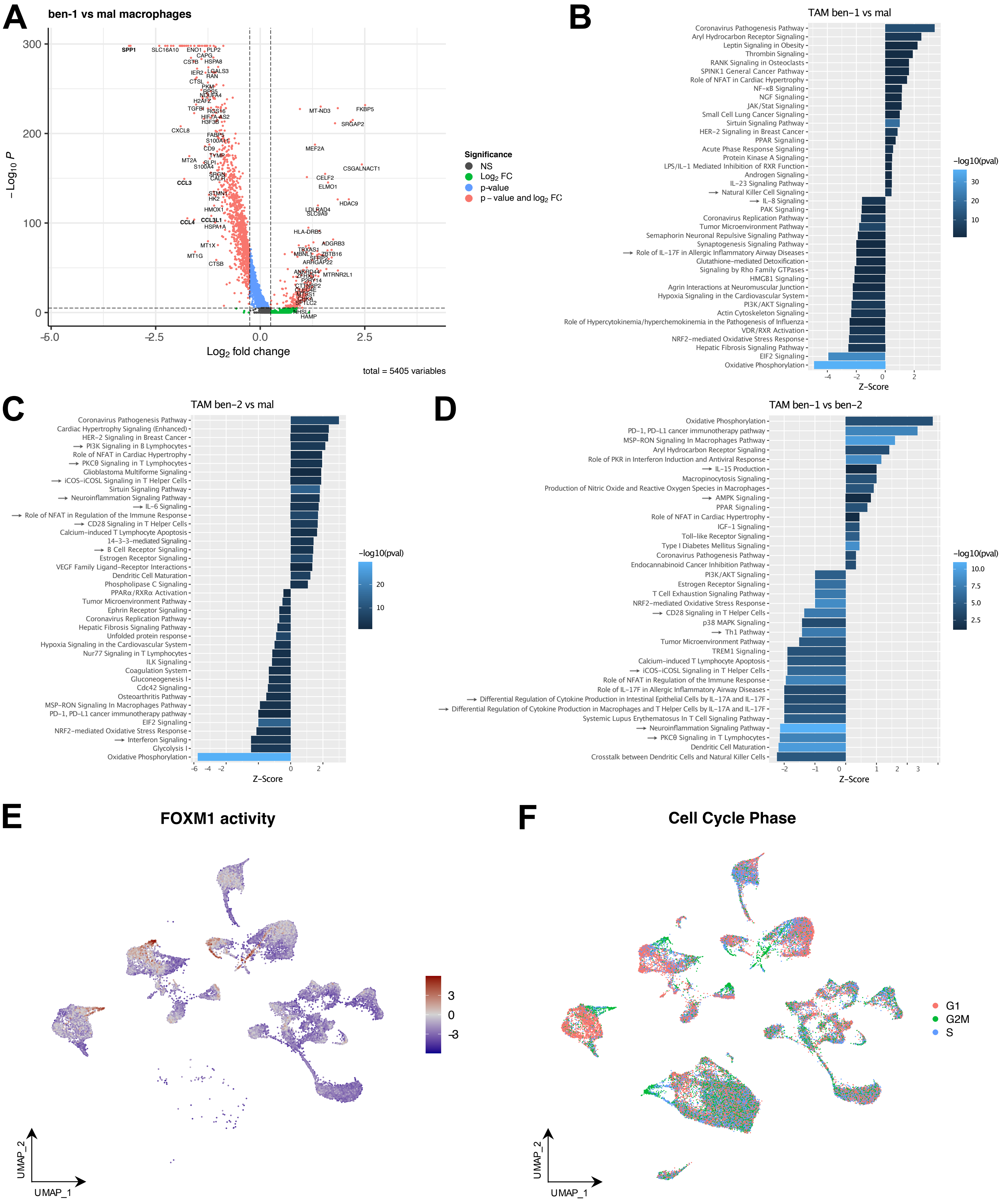

### Suppl. Fig. 5

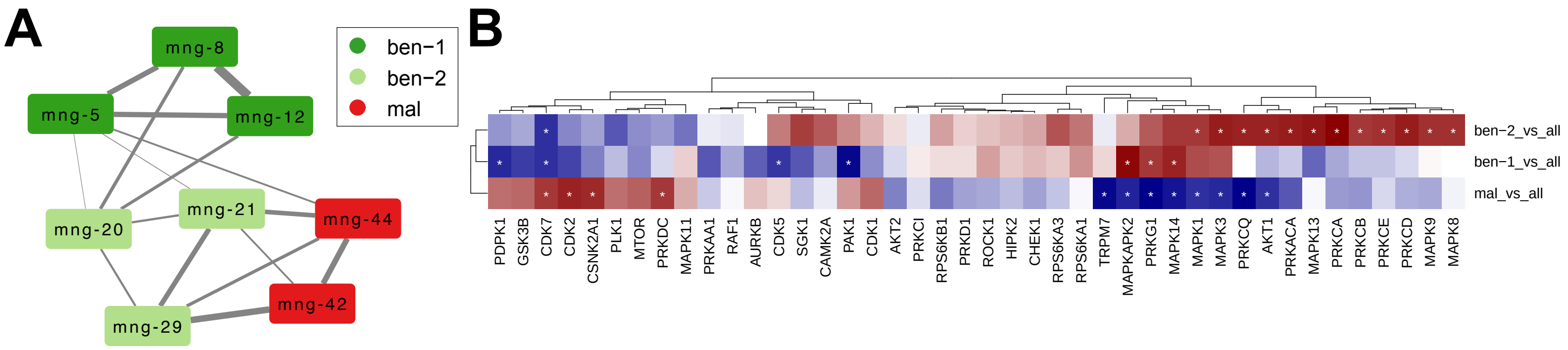
