## Supplementary material for "Integrated phospho-proteogenomic and single-cell transcriptomic analysis of meningiomas establishes robust subtyping and reveals subtype-specific immune invasion": Suppl. Fig. 4

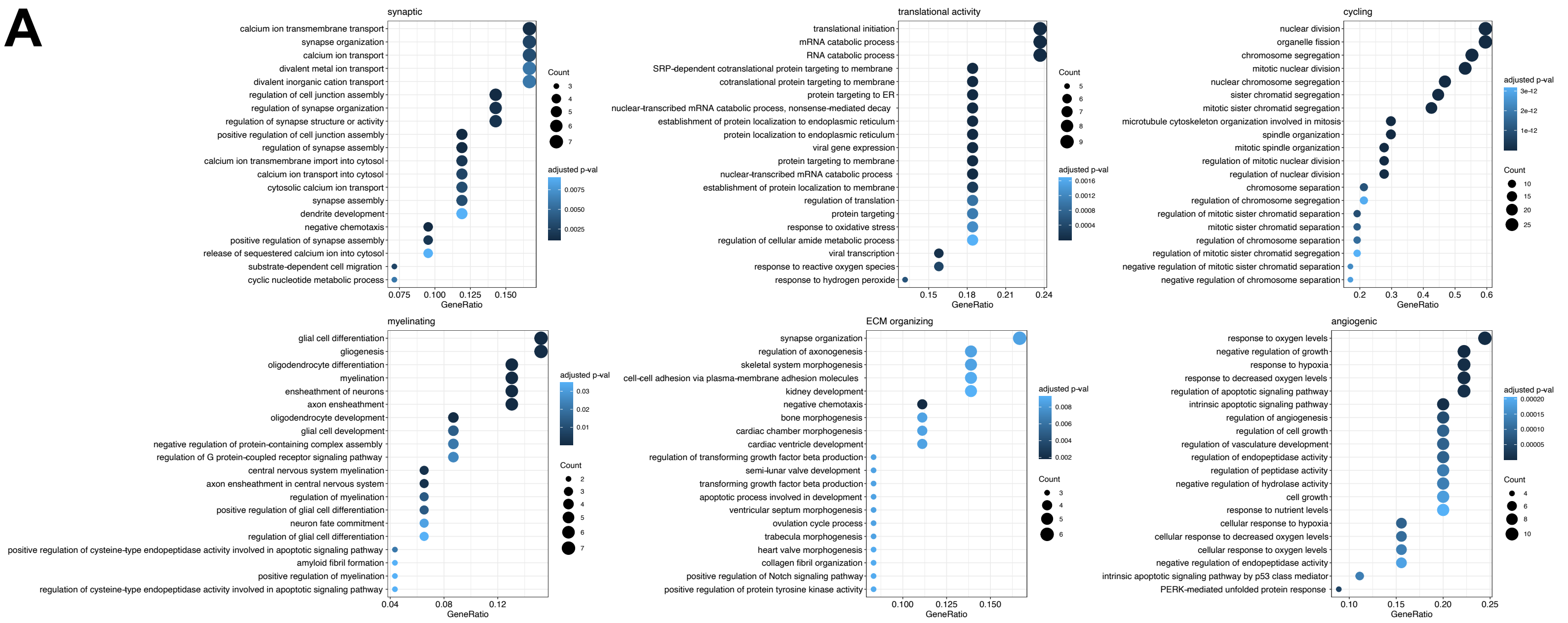


**B**

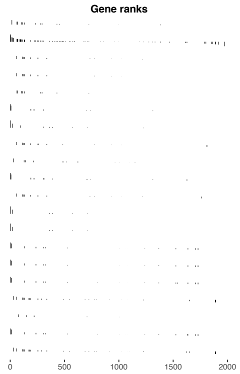

| Pathway | NES | pval | padj |
|---|---|---|---|
| Protein-protein interactions at synapses | 2.01 | 1.9e-04 | 2.6e-03 |
| Neuronal System | 1.88 | 2.5e-04 | 2.7e-03 |
| → PI5P, PP2A and IER3 Regulate PI3K/AKT Signaling | 1.85 | 2.3e-03 | 1.4e-02 |
| → Constitutive Signaling by Aberrant PI3K in Cancer | 1.85 | 1.4e-03 | 1.1e-02 |
| Neurexins and neuroligins | 1.81 | 1.9e-03 | 1.2e-02 |
| Nitric oxide stimulates guanylate cyclase | 1.73 | 8.8e-03 | 4.0e-02 |
| GABA receptor activation | 1.71 | 8.3e-03 | 3.9e-02 |
| → PI3K/AKT Signaling in Cancer | 1.67 | 9.1e-03 | 4.1e-02 |
| Molecules associated with elastic fibres | 1.67 | 2.2e-02 | 8.8e-02 |
| Platelet homeostasis | 1.66 | 2.0e-02 | 8.0e-02 |
| → Negative regulation of the PI3K/AKT network | 1.65 | 1.1e-02 | 4.9e-02 |
| Activation of GABAB receptors | 1.63 | 1.7e-02 | 7.0e-02 |
| GABA B receptor activation | 1.63 | 1.7e-02 | 7.0e-02 |
| Defective B3GALTL causes Peters-plus syndrome (PpS) | 1.62 | 3.6e-02 | 1.1e-01 |
| O-glycosylation of TSR domain-containing proteins | 1.62 | 3.6e-02 | 1.1e-01 |
| Diseases associated with O-glycosylation of proteins | 1.60 | 2.3e-02 | 8.8e-02 |
| → RAF/MAP kinase cascade | 1.60 | 2.6e-02 | 9.8e-02 |
| Netrin-1 signaling | 1.59 | 3.5e-02 | 1.1e-01 |
| O-linked glycosylation | 1.58 | 2.7e-02 | 9.9e-02 |
| → MAPK1/MAPK3 signaling | 1.55 | 3.8e-02 | 1.2e-01 |

**C**

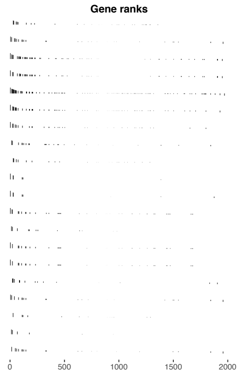

| Pathway | NES | pval | padj |
|---|---|---|---|
| Chemokine receptors bind chemokines | 1.93 | 1.3e-04 | 7.2e-03 |
| Collagen formation | 1.91 | 2.4e-04 | 1.1e-02 |
| Cellular responses to external stimuli | 1.91 | 2.0e-05 | 3.3e-03 |
| Cellular responses to stress | 1.87 | 8.6e-05 | 5.4e-03 |
| Cytokine Signaling in Immune system | 1.82 | 1.5e-05 | 3.3e-03 |
| Signaling by Interleukins | 1.82 | 5.3e-05 | 3.8e-03 |
| Interleukin-4 and Interleukin-13 signaling | 1.81 | 6.4e-04 | 2.5e-02 |
| ECM proteoglycans | 1.80 | 1.3e-03 | 3.5e-02 |
| Interleukin-10 signaling | 1.79 | 1.4e-03 | 3.5e-02 |
| → VEGFA-VEGFR2 Pathway | 1.78 | 1.4e-04 | 7.3e-03 |
| → Signaling by VEGF | 1.78 | 3.1e-04 | 1.3e-02 |
| Platelet activation, signaling and aggregation | 1.78 | 9.6e-04 | 3.5e-02 |
| → Signaling by TGF-beta family members | 1.77 | 1.0e-03 | 3.5e-02 |
| Platelet degranulation | 1.76 | 1.3e-03 | 3.5e-02 |
| Response to elevated platelet cytosolic Ca2+ | 1.76 | 1.3e-03 | 3.5e-02 |
| FOXO-mediated transcription | 1.74 | 2.6e-03 | 4.9e-02 |
| Assembly of collagen fibrils and other multimeric structures | 1.73 | 2.1e-03 | 4.6e-02 |
| Extra-nuclear estrogen signaling | 1.72 | 3.1e-03 | 5.4e-02 |
| → Signaling by TGF-beta Receptor Complex | 1.71 | 1.3e-03 | 3.5e-02 |
| Collagen biosynthesis and modifying enzymes | 1.71 | 5.5e-03 | 6.5e-02 |
